# Gene identity, not variant effect, dominates ClinVar benchmarks of missense pathogenicity predictors

**DOI:** 10.64898/2026.08.21.746180

**Authors:** Saad Harrizi, Imane Nait Irahal, Kabine Mostafa, Damien Arnoult

## Abstract

Missense pathogenicity predictors are routinely benchmarked against ClinVar, whose labels are strongly structured by gene: genes under diagnostic scrutiny accumulate pathogenic submissions while incidentally sequenced genes accumulate benign ones. We asked how much of a benchmark score this structure alone can produce. On 197,904 ClinVar missense variants validated against UniProt canonical sequences, a null model using no variant-level information, scoring each variant only by the pathogenic fraction of its own gene, reaches an area under the receiver operating characteristic curve (AUROC) of 0.921 under a random 10-fold split. On a common intersection of 169,989 variants, four current predictors exceed it by only 0.036 to 0.044. The inflation is not uniform, so it does not cancel when predictors are compared: under within-gene evaluation the ranking inverts, AlphaMissense rising from third to first and gMVP falling to third (p < 0.0001). The inversion survives removal of ceiling genes and replicates on an independently curated benchmark. Because both rankings derive from the same ClinVar labels, we arbitrated between them using data with no gene-level structure: agreement with 47 human deep mutational scanning assays matches the within-gene ranking and inverts the conventional one (p = 0.027, 0.0023). Across twenty-two dbNSFP predictors scored on one common intersection of 112,248 variants, with each tool’s exposure to clinical labels registered before any score was extracted, predictors never trained on such labels sit 0.051 AUROC behind supervised ones globally but only 0.026 behind within genes (difference +0.025 [+0.023, +0.027], p < 0.0001). Leave-one-out correction, the standard remedy, is worth 0.002 AUROC. Much of ClinVar benchmark performance reflects gene identity rather than variant effect, and the distortion changes which predictor a benchmark ranks first, in a direction experimental data contradicts. We release genenull, a single-file implementation, so reporting this baseline costs one function call.

**Author summary:** When a computer program predicts whether a genetic variant causes disease, we judge it by testing it against ClinVar, a public archive of variants clinicians have already interpreted. We found that this test is easier to pass than it looks. Some genes appear in ClinVar because they are suspected of causing disease, so most of their recorded variants are harmful; others are sequenced incidentally, so most of theirs are harmless. A program that knows nothing about a variant except which gene it sits in can exploit that pattern, and scores almost as well as the best tools available. This matters beyond a single number. When we removed the gene pattern and ranked variants inside a single gene, the order changed: the tool that looked best became worst, and the one that looked worst became best. Laboratory experiments that measure the effect of every possible variant in a protein agree with the new order, not the old one. Across twenty-two prediction tools, about half the advantage held by programs trained on clinical data disappears once the gene pattern is removed. We release software so anyone can measure this baseline in one line of code.

## 1. Introduction

Interpreting missense variants is a central problem in clinical genomics, and a large family of computational predictors now addresses it, spanning conservation and structure-based methods [1, 2], ensembles and genome-wide annotation scores [3, 4], supervised deep networks [5], unsupervised generative models [6], and proteome-wide predictors trained at scale [7]. Evaluation of these tools rests overwhelmingly on ClinVar, the public archive of clinically interpreted variants [8]. Reported areas under the ROC curve on ClinVar are commonly in the 0.90–0.95 range [9, 10, 11] and are used to rank tools against one another, to justify deployment in diagnostic pipelines, and to motivate new specialised models.

Two properties of ClinVar complicate this. First, submissions are not distributed uniformly across the genome: they concentrate in genes with established disease associations, and within those genes, ascertainment is driven by clinical testing panels. Second, the label a variant receives is correlated with the gene it sits in. Genes under diagnostic scrutiny for a dominant disease accumulate pathogenic entries; genes sequenced incidentally accumulate benign ones.

*Relation to what is already known.* That ClinVar-based evaluation is gene-dependent is not news, and we want to be precise about what is and is not new here. Circularity in variant effect prediction was formalised by Grimm and colleagues as type 1, where the same variants appear in training and test data, and type 2, where variants from the same protein do [12]. More recently, Tejura and colleagues showed that calibrations derived genome-wide do not transfer to individual genes: predicted score distributions are not uniform across genes, around 70% of evaluable evidence-strength intervals were discordant, and many variants would receive inappropriately strong or weak evidence under a genome-wide calibration [13]. Related work documents that per-gene AUROC varies with gene function, structure and conservation across dozens of predictors [14], and large benchmarking efforts have flagged that training-data circularity is hard to detect when models were fitted to historical ClinVar releases [9]. ClinGen’s recommendations for PP3/BP4 evidence thresholds, which sit within the American College of Medical Genetics and Genomics and Association for Molecular Pathology (ACMG/AMP) interpretation framework, rest on genome-wide calibration of exactly this kind [15, 16].

That literature concerns calibration: whether a given score corresponds to the same strength of evidence in every gene. Our question is about discrimination: how much of the ranking performance a benchmark reports is available without looking at the variant at all. The two are distinct, and neither implies the other. A predictor could be perfectly calibrated in every gene and still owe most of its AUROC to gene identity; it could equally be poorly calibrated while discriminating on genuine variant-level signal.

Two studies come close. Grimm et al. named this structure type 2 circularity and treated it as contamination between training and test sets [12]. Lin et al. observed that many ClinVar genes carry variants of a single class, reasoned that a model could therefore learn the gene rather than the variant, and tested it by rebuilding a label-balanced subset on which their predictor reaches AUROC 0.88 [17]. Neither measures what gene identity is worth on its own, and to our knowledge no previous study constructs the corresponding null. We do so directly: a model given nothing but the gene label, no sequence, no structure, no conservation, no information about which substitution occurred, reaches AUROC 0.921, against 0.955 to 0.963 for four current predictors on the same variants. Prior work establishes that predictor accuracy *varies* by gene [13, 14]; we show that under the split scheme conventionally used to report benchmark performance, gene identity alone reproduces most of the reported score. The first finding says a benchmark score is an average over unequal parts, and is addressed by calibrating per gene. The second says most of that score does not require the variant, and is not addressed that way at all.

We are deliberate about the word *reproduces*. The null is not a predictor and we never claim it is: under leave-gene-out splitting it falls to exactly 0.500, since it can say nothing about a gene it has not seen. The claim is about what the benchmark measures, not about what the null could be used for.

We also show that the standard remedy is inadequate rather than merely imperfect. Supervised predictors ship leave-one-out scores precisely to address training-set contamination. Leave-one-out removes the variant itself but leaves every other variant in its gene in training, so the gene-level structure survives untouched: the correction is worth 0.002 AUROC against an effect of 0.036 to 0.044. The confound therefore survives elimination of both type 1 and type 2 circularity, because no variant and no protein need be shared for it to operate, it is a property of how the archive was assembled rather than of any particular split.

The practical consequence concerns specialisation. A recurring pattern in the literature is to select a gene set of biological interest, benchmark general predictors on it, observe that performance differs from the genome-wide average, and conclude that the gene set requires dedicated modelling. If gene-level label structure varies between gene sets, and it does, then this inference is unsafe. We demonstrate the point on mitochondrial proteins, where we ourselves pursued exactly this reasoning in an earlier and unpublished analysis that this work supersedes.

## 2. Results

### 2.1 A gene-identity null model approaches state-of-the-art performance

We assembled 197,904 ClinVar missense variants with unambiguous pathogenic or benign classifications, each validated against the UniProt canonical sequence for its gene [18] (Fig 1a; Methods, “Variant dataset”). AlphaMissense scores were available for 195,270 of these (98.7%), joined on UniProt accession and protein-level substitution so that no genomic liftover was required.

**Fig 1.**
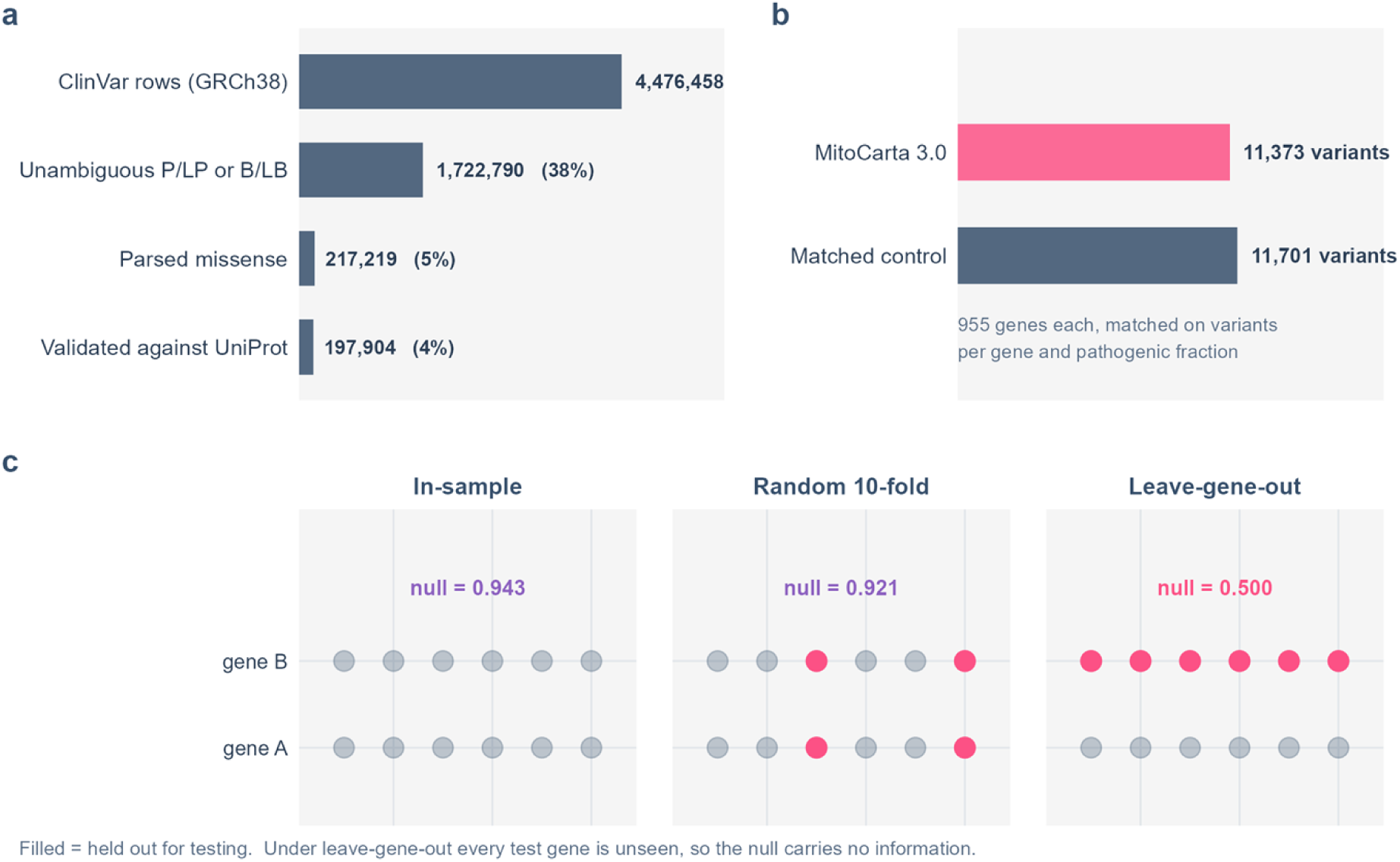
Study design. **a**, Attrition from the raw ClinVar release to the analysed set, drawn so that bar length is the count: 4,476,458 rows reduce to 197,904 missense variants validated against UniProt canonical sequences. **b**, The mitochondrial set and its non-mitochondrial control, matched gene-by-gene on variants per gene and pathogenic fraction, 955 genes each. **c**, What each split scheme does to the gene-identity null. Filled marks are held out for testing; under leave-gene-out every test gene is unseen, so the null carries no information and falls to 0.500.

The null model assigns each variant the fraction of variants in its own gene that are labelled pathogenic. Evaluated in-sample it reaches AUROC 0.943 [0.938, 0.948]. Evaluated under a random 10-fold variant-level split, the scheme used throughout much of the literature, in which the same gene appears in both training and test folds, it reaches 0.921 [0.915, 0.927]. AlphaMissense on the same variants reaches 0.956. Under leave-gene-out splitting the null falls to exactly 0.500, because every held-out gene is unseen and receives an identical constant (Fig 2a).

**Fig 2.**
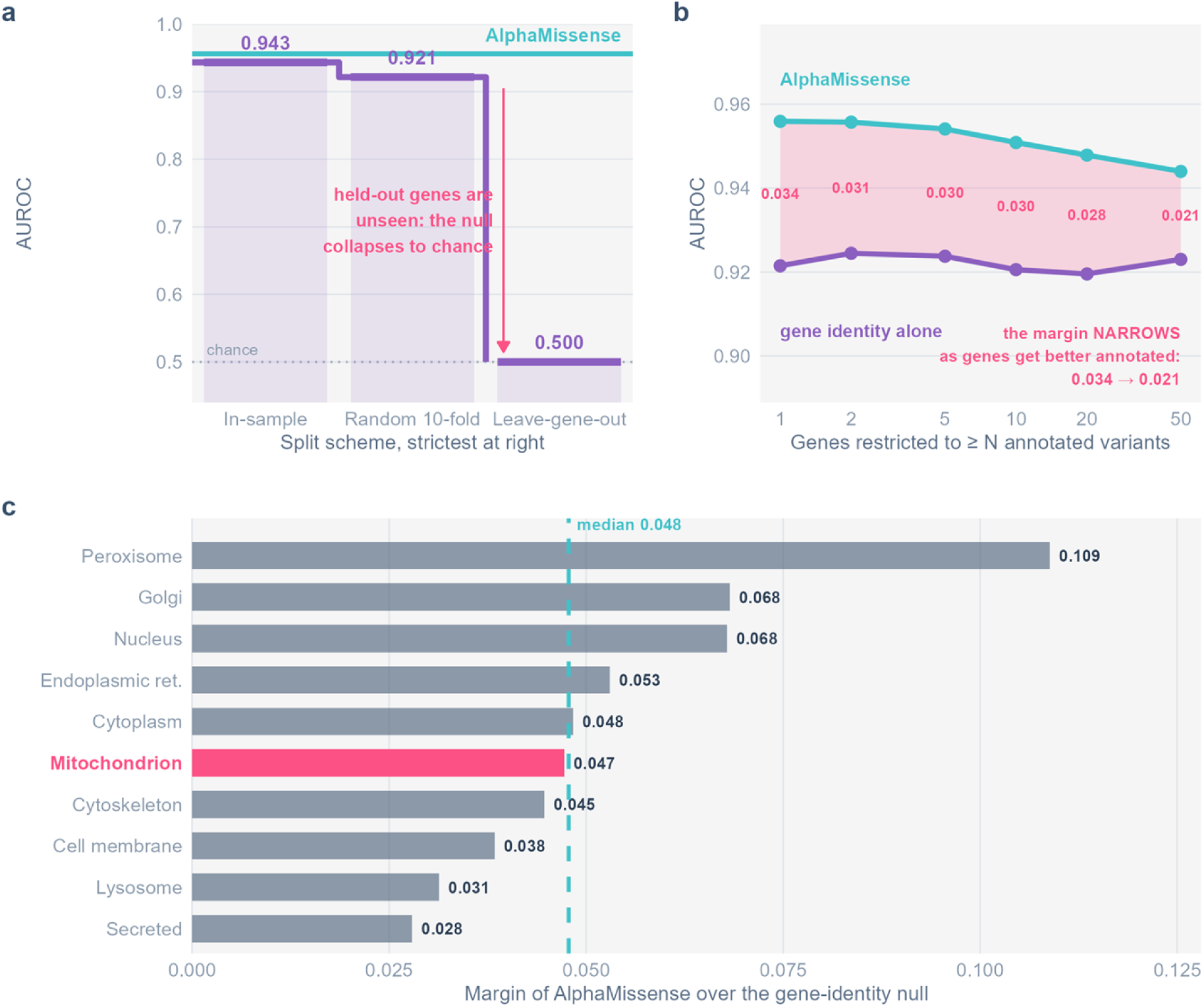
The gene-identity null. **a**, The null under three split schemes against AlphaMissense on the same variants. **b**, Robustness to gene annotation depth: as the minimum variants per gene rises from 1 to 50, the fraction of label-pure genes falls almost tenfold while the null’s AUROC does not move. **c**, Predictor and null in each of ten subcellular compartments, with the margin between them.

The margin between a state-of-the-art predictor and a model that knows only which gene a variant is in is therefore 0.035 [0.019, 0.051] under the split scheme most commonly used to report benchmark performance.

*This is not specific to AlphaMissense.* We repeated the comparison across four current predictors on a single common intersection of 169,989 variants in 13,195 genes, so that every method is evaluated on identical data rather than on its own coverage subset (Fig 3). Against the gene-identity null’s 0.9191 on this intersection, VARITY_R [19] reaches 0.9635, VARITY_R_LOO 0.9615, gMVP [20] 0.9590 and AlphaMissense 0.9548 (S1 Table).

**Fig 3.**
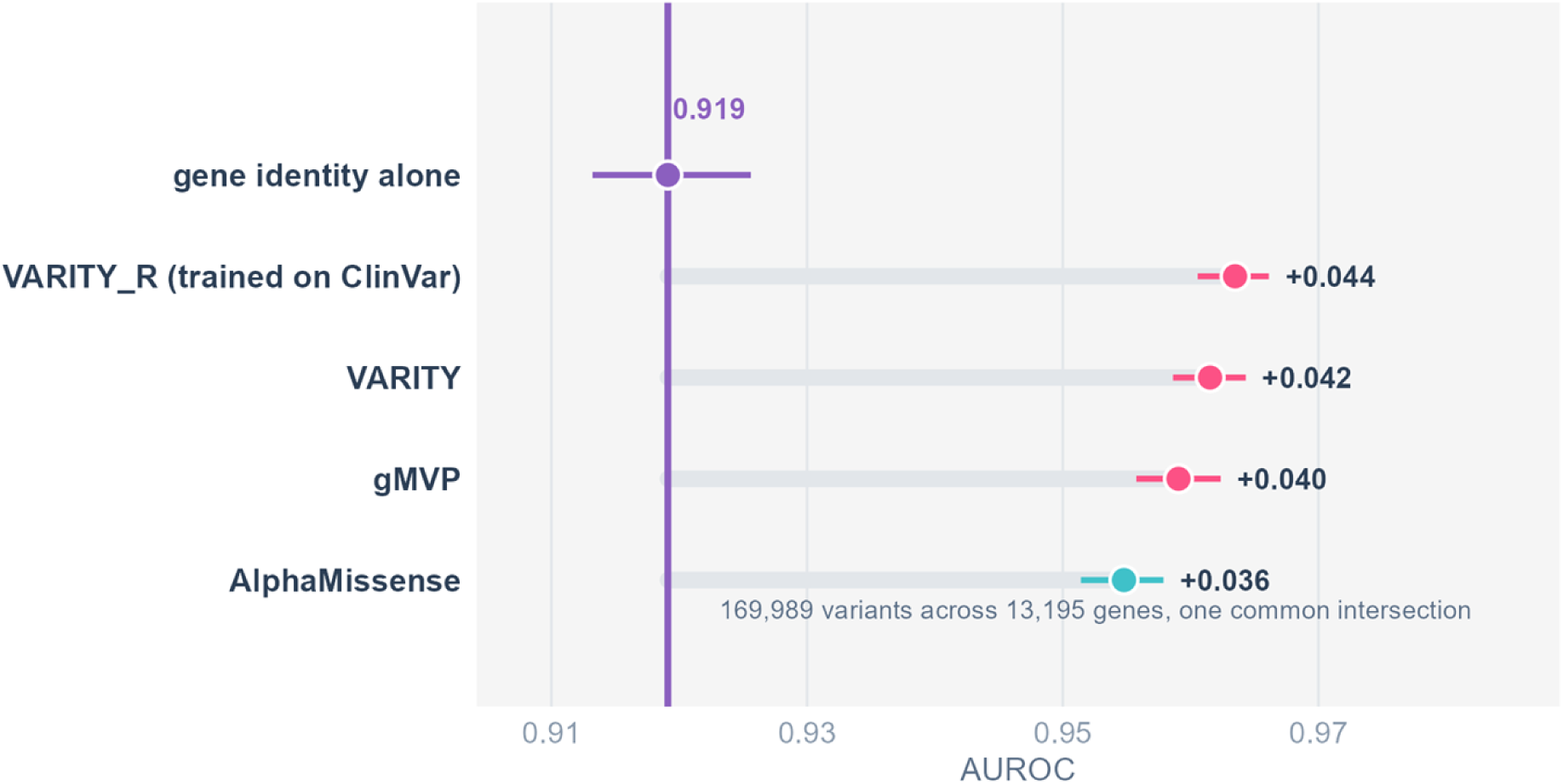
All comparators against the gene-identity null on a single common intersection of 169,989 variants in 13,195 genes, so that every method is scored on identical data rather than on its own coverage subset.

Every method lies between 0.036 and 0.044 above the null. The spread among the four predictors is 0.009 AUROC, roughly a fifth of the distance any of them stands above a model with no variant-level information at all.

A 0.009 difference on 169,989 variants is far outside sampling error, so choosing between tools on that basis is reasonable. The point is that the comparison is made on a narrow and contaminated margin and, as the next section shows, the ordering on that margin is not the one that survives removing the contamination.

### 2.2 The field’s own correction addresses two thousandths of the effect

The standard remedy for training-set contamination is the leave-one-out score, which supervised predictors ship for exactly this purpose. VARITY is trained on rare ClinVar variants and releases a leave-one-out column in which each training variant is excluded in turn and predicted by a model that did not see it [19]. Its authors also excluded features informed by protein identity for exactly this reason, which makes it the strongest available test: the gene structure we describe survives both precautions, because it is carried by the labels rather than by the features. Applied as its authors intended, that correction costs 0.002 AUROC (0.9635 → 0.9615), against a gene-identity null worth 0.919 and a margin over that null of 0.036 to 0.044.

The reason is structural rather than incidental. Leave-one-out removes the variant itself but leaves every *other* variant in the same gene in the training set, so the gene-level label structure survives it intact. Type 1 circularity, in the established terminology, is worth two thousandths here; the gene-level structure that no leave-one-out scheme touches is worth two orders of magnitude more. This is the clearest evidence that the confound we describe is not the one the field has already addressed, and it is not a matter of degree: no variant-level correction can reach a gene-level structure.

### 2.3 Gene-controlled evaluation reverses which predictor wins

If every predictor were inflated by the same amount, relative comparisons would survive and the practical consequence of the above would be limited, benchmarks are used mainly to rank tools against one another. We tested this directly.

Within-gene AUROC is the natural gene-controlled measure of discrimination. It asks whether a predictor ranks variants correctly *inside* a single gene, where between-gene label structure contributes nothing by construction: every comparison is between two variants sharing a gene, so the gene prior is constant and carries no information. Global AUROC mixes this ability with the separate ability to separate pathogenic genes from benign ones; within-gene AUROC isolates it.

We do not claim within-gene AUROC is *the* clinically relevant metric; a clinician needs an absolute classification, for which calibration matters more. The claim is about measurement: within-gene AUROC is what global AUROC would equal if gene-level structure carried no information, so comparing the two isolates how much of a benchmark score depends on it.

Within-gene AUROC is only defined for genes carrying both labels, so this analysis is restricted to the 1,860 genes with at least three variants of each class. That is 12.5% of genes but 48.6% of variants (96,142), and those genes are by construction the better-characterised ones. The restriction is unavoidable, the quantity does not exist elsewhere, but it means the reversal is established on well-annotated genes and is not demonstrated for sparsely annotated ones. On that set the ranking inverts (Fig 4a; Table 1):

**Fig 4.**
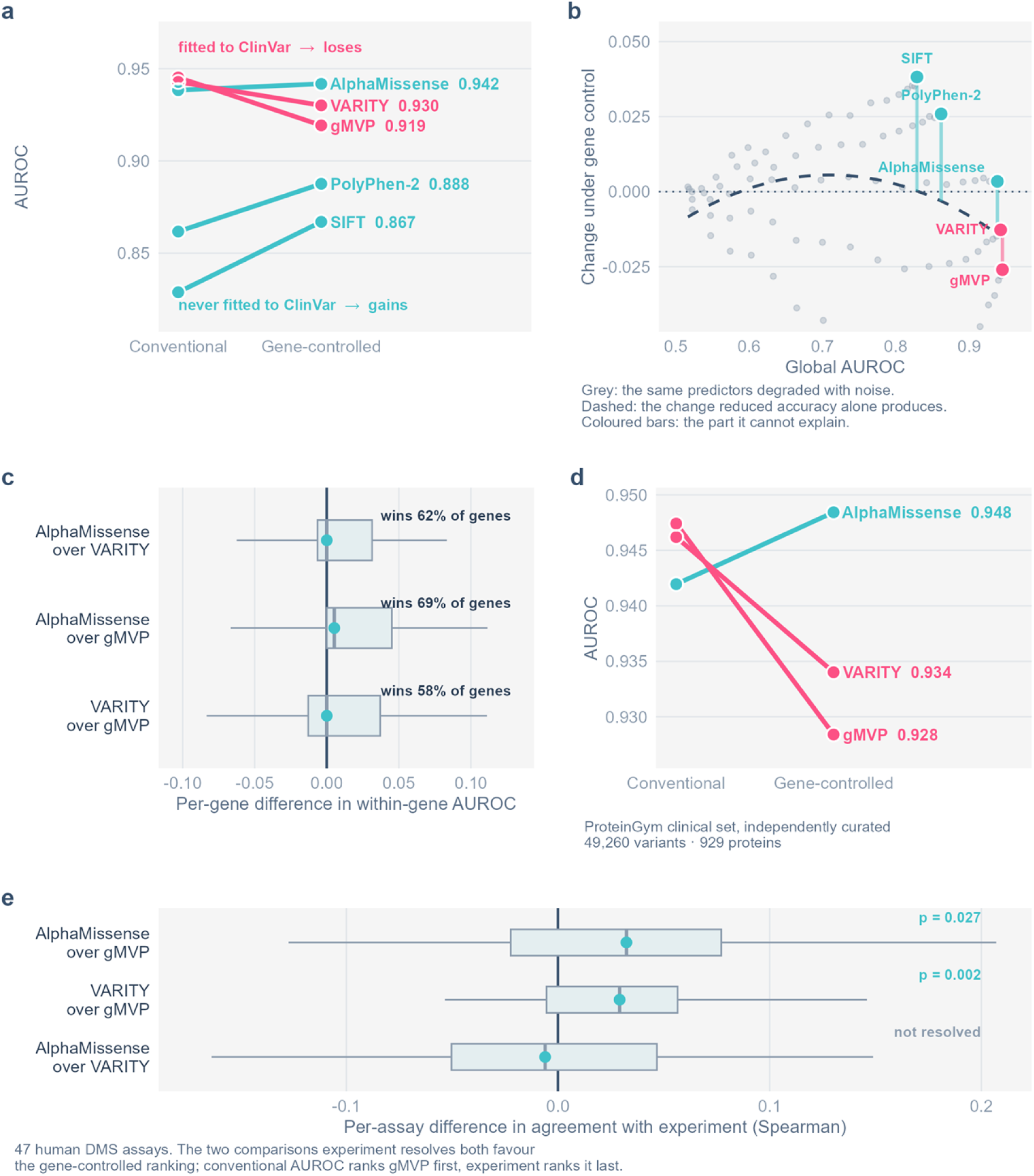
Gene-controlled evaluation reverses the ranking. **a**, Slopegraph of global to within-gene AUROC across five predictors spanning two decades; predictors never fitted to ClinVar gain, those supervised on it lose. **b**, The headroom control. Because the change in **a** is perfectly rank-correlated with global AUROC, the same picture is predicted by “weaker predictors have more room to move”. Grey marks are those five predictors degraded with uniform noise, which lowers global AUROC while changing nothing about their training; the dashed line is fitted to them and is what headroom alone produces. Coloured bars are the residuals, the part headroom cannot explain, spanning 4.1 times the range the null curve covers. **c**, Per-gene paired differences with win rates from Wilcoxon signed-rank tests. **d**, Independent replication on the ProteinGym clinical substitutions benchmark, curated by a different group through a different pipeline. **e**, Per-assay differences in agreement with experiment across 47 human ProteinGym deep mutational scanning assays. Both comparisons that experiment resolves favour the gene-controlled ranking; conventional AUROC ranks gMVP first, experiment ranks it last.

**Table 1.** Global and within-gene AUROC for the three modern predictors.

| Predictor | Global AUROC | Rank | Within-gene AUROC | Rank |
| --- | --- | --- | --- | --- |
| VARITY_R_LOO | 0.9615 | 1 | 0.9306 | 2 |
| gMVP | 0.9590 | 2 | 0.9114 | 3 |
| AlphaMissense | 0.9548 | 3 | 0.9381 | 1 |

*Note.* Global AUROC is computed on the full common intersection of 169,989 variants; within-gene AUROC on the 1,860 genes carrying at least three variants of each class (96,142 variants), where the quantity is defined.

AlphaMissense goes from last to first. Two of three pairwise comparisons reverse sign, both decisively: VARITY minus AlphaMissense moves from +0.0067 globally to −0.0075 [−0.0106, −0.0045] within genes, and AlphaMissense minus gMVP from −0.0043 to +0.0267 [+0.0233, +0.0301], each at p < 0.0001. VARITY minus gMVP does not change sign but widens from +0.0025 to +0.0192 [+0.0156, +0.0229] (S2 Table).

*The reversal is robust to every check we applied.* It is not an artefact of how genes are aggregated: AlphaMissense ranks first under an unweighted mean over genes, a variant-weighted mean, and the median. Beyond that:

- *Threshold sensitivity.* The evaluable gene set depends on how many variants of each class a gene must carry. The ordering is identical at every threshold tested, at least 2, 3, 5, 10 or 20 of each class, spanning 2,440 genes down to 214. This is the most informative of the three checks, because the 214-gene set is both small and differently composed.
- *Split-half and leave-one-gene-out.* The ordering is unchanged across 200 random half-partitions and across removal of any single gene. Both are weak tests at this sample size and would only have been informative had they failed.

*Ceiling effects and ties.* A substantial minority of genes are perfectly separated: 533, 499 and 408 of the 1,860 genes reach AUROC exactly 1.0 for AlphaMissense, VARITY and gMVP respectively, and in 558 genes (30.0%) two or more predictors tie at the ceiling. This compresses the mean and makes naive per-gene win counts meaningless, assigning tied genes by column order produces a different answer depending on the order chosen. Counting only the 1,302 genes with a unique best predictor, AlphaMissense wins 600 (46.1%), VARITY 457 (35.1%) and gMVP 245 (18.8%), consistent with the ordering by mean.

The ordering survives removal of the ceiling entirely: it is unchanged when genes where any predictor reaches 1.0 are excluded (n = 1,123) and when genes where all three reach 1.0 are excluded (n = 1,608).

Because the mean is a questionable summary in the presence of a ceiling, we also report paired rank-based tests, which make no assumption about the distribution of per-gene values (Fig 4c; S3 Table). AlphaMissense wins 70.0% of genes against gMVP (1,035/1,479, p = 7×10⁻⁵⁸), VARITY 64.7% against gMVP (962/1,488, p = 6×10⁻³⁰), and AlphaMissense 56.8% against VARITY (804/1,415, p = 1×10⁻⁶).

All three agree with the mean-based ordering. The AlphaMissense advantage over VARITY is the smallest of the three, a 57% win rate rather than 70%, and we present it as such rather than as a decisive separation.

*Why this pattern is expected under our hypothesis, and one complication.* VARITY and gMVP are both supervised on clinical labels, VARITY on rare ClinVar variants and gMVP on ClinVar together with HGMD, UniProt and SwissVar [19, 20]. AlphaMissense is not fitted to clinical labels at all; ClinVar enters only when its thresholds are calibrated to 90% precision [7]. Under conventional evaluation the two supervised models lead; once between-gene label structure is removed, the model that never fitted those labels leads instead. This is what our thesis predicts: supervised training on a gene-structured archive buys performance that does not survive gene control. Three predictors are too few to establish that on their own, which is why the next section tests it across twenty-two.

*A dose–response across twenty years of methodology.* To test the mechanism against a wider range of ClinVar exposure we added PolyPhen-2 (2010) [2] and SIFT (2003) [1], obtained from the EBI Proteins API [21]. Both predate modern ClinVar; SIFT uses sequence conservation alone and was never trained on clinical labels at all. On the five-predictor intersection (59,045 variants, 1,055 evaluable genes), the change from global to within-gene AUROC orders monotonically by exposure to ClinVar (S11 Table): Every predictor trained on ClinVar loses under gene control; every predictor that was not, gains. ESM-2 zero-shot [22], which has seen no clinical label of any kind, gains +0.0167 and fits the same ordering. The gradient spans five tools built on entirely different principles across two decades, and it is monotone.

*The gradient is confounded with global performance, so we tested it against a headroom null.* This ordering is also, exactly, the ordering by global AUROC: across the five predictors the rank correlation between global AUROC and the change is −1.000. A weaker predictor has more room to move, so regression to the mean predicts the same gradient with no mechanism at all, and five points cannot separate the two explanations by inspection. We therefore constructed the null directly. Each predictor was degraded by mixing its rank-scaled score with uniform noise, in twenty steps, and the within-gene change was recomputed at each step (Fig 4b; Methods, “Headroom control”). Degradation changes nothing about what a predictor was trained on; it only lowers global AUROC. The resulting curve is what headroom alone produces.

Headroom alone produces very little. Across the full range of global AUROC spanned by the five predictors, the null curve moves by −0.0155, whereas the observed changes span 0.0642, 4.1 times as much, and in the opposite direction at the low end. At SIFT’s global AUROC the null predicts a change of +0.0002; SIFT gains +0.0382. Expressed as a share, degradation accounts for 24% of the observed spread and none of the gains (S12 Table).

### 2.4 The gradient across twenty-two predictors

We extracted all 37 per-predictor score columns from dbNSFP 5.3.1a [23] for our variants. Each predictor’s exposure to clinical labels was classified from its published training description and recorded before any dbNSFP score was extracted (S1 Text), on three levels: no clinical labels at any stage (0), labels used for calibration or a non-clinical proxy target (1), and supervision directly on curated clinical labels (2). Predictors taking other listed predictors as input features were excluded as non-independent, and where a model contributes several columns (dbNSFP ships four VARITY variants, three Meta*, and two each of PolyPhen-2, SIFT, MisFit and BayesDel), a single pre-specified representative was used, so no model is counted more than once.

Twenty-two representatives cover at least half our variants. Their common intersection is 112,248 variants in 10,458 genes, and every value below is computed on exactly those rows (Fig 5a; Table 2; Methods, “The dbNSFP panel”).

**Fig 5.**
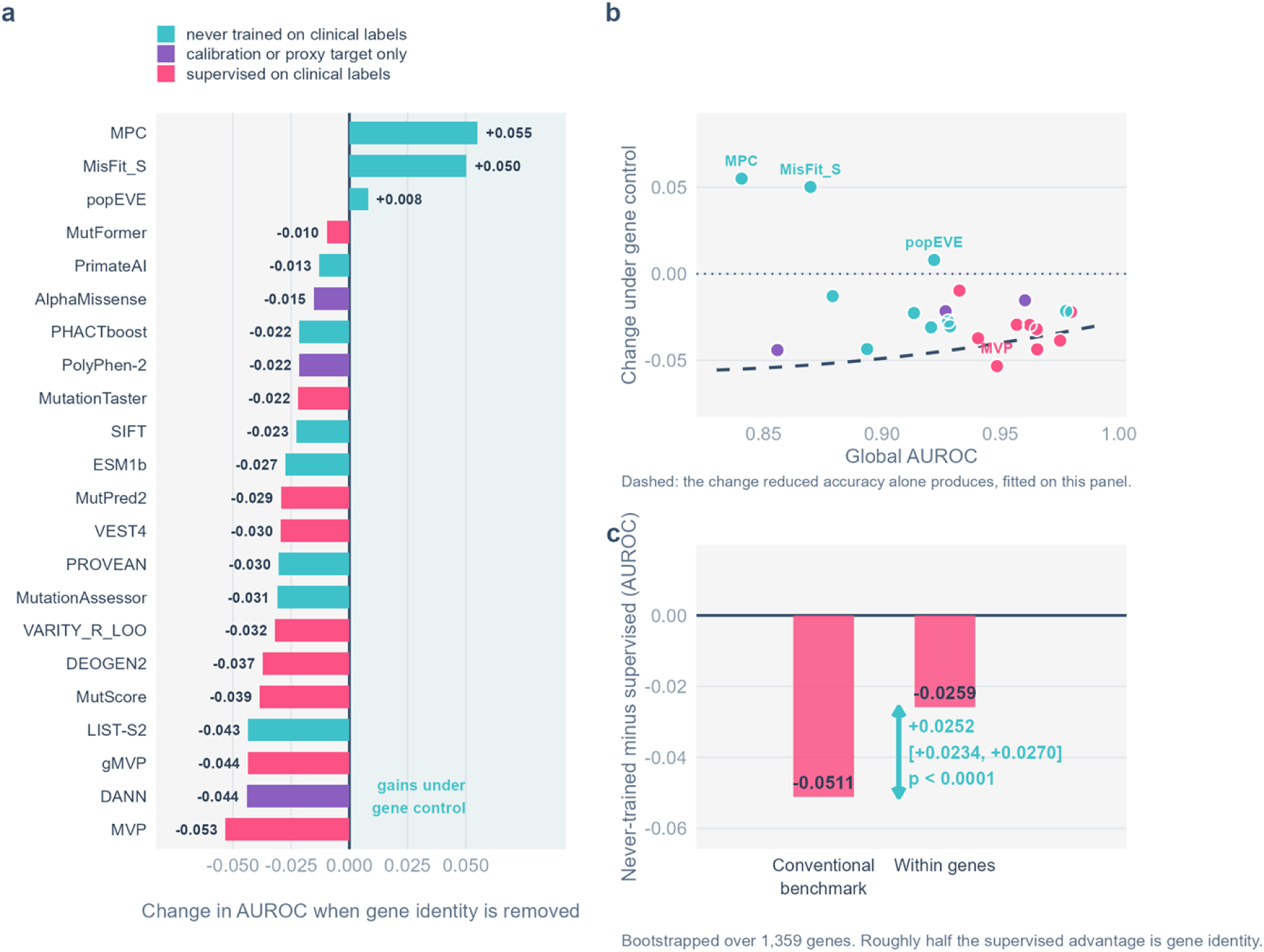
The exposure gradient across the dbNSFP panel. **a**, Change in AUROC under gene control for each of twenty-two predictors on their common intersection, ordered, coloured by exposure to clinical labels as pre-registered before any score was extracted. The three predictors that gain are the three whose training signal is population genetics or evolution rather than annotation. **b**, The same predictors against the headroom null fitted on this panel’s own intersection; the dashed line is the change that reduced accuracy alone produces. **c**, The never-trained group’s deficit against the supervised group, measured on a conventional benchmark and again within genes. The gap between the two bars is the difference-in-differences, bootstrapped over 1,359 genes.

**Table 2.** Change from global to within-gene AUROC across twenty-two predictors on one common intersection of 112,248 variants in 10,458 genes.

| Predictor | Exposure | Global | Within-gene | Change |
| --- | --- | --- | --- | --- |
| MPC | 0 none | 0.8408 | 0.8958 | +0.0550 |
| MisFit_S | 0 none | 0.8699 | 0.9201 | +0.0502 |
| popEVE | 0 none | 0.9220 | 0.9300 | +0.0080 |
| MutFormer | 2 supervised | 0.9327 | 0.9230 | -0.0097 |
| PrimateAI | 0 none | 0.8792 | 0.8663 | -0.0129 |
| AlphaMissense | 1 calibration only | 0.9604 | 0.9451 | -0.0153 |
| PHACTboost | 0 none | 0.9777 | 0.9562 | -0.0215 |
| PolyPhen-2<br>(HVAR) | 1 calibration only | 0.9269 | 0.9053 | -0.0216 |
| MutationTaster | 2 supervised | 0.9798 | 0.9578 | -0.0220 |
| SIFT | 0 none | 0.9135 | 0.8909 | -0.0226 |
| ESM1b | 0 none | 0.9280 | 0.9005 | -0.0275 |
| MutPred2 | 2 supervised | 0.9569 | 0.9275 | -0.0294 |
| VEST4 | 2 supervised | 0.9624 | 0.9328 | -0.0295 |
| PROVEAN | 0 none | 0.9287 | 0.8984 | -0.0303 |
| MutationAssessor | 0 none | 0.9207 | 0.8898 | -0.0309 |
| VARITY_R_LOO | 2 supervised | 0.9653 | 0.9333 | -0.0320 |
| DEOGEN2 | 2 supervised | 0.9406 | 0.9033 | -0.0372 |
| MutScore | 2 supervised | 0.9752 | 0.9366 | -0.0386 |
| LIST-S2 | 0 none | 0.8938 | 0.8503 | -0.0435 |
| gMVP | 2 supervised | 0.9655 | 0.9219 | -0.0436 |
| DANN | 1 calibration only | 0.8559 | 0.8119 | -0.0441 |
| MVP | 2 supervised | 0.9486 | 0.8952 | -0.0533 |

*Note.* Exposure to clinical labels was pre-registered before any score was extracted.

*The gradient is noisier across twenty-two tools than across five, as it should be*. The five-predictor ordering was perfectly monotone; across this panel the rank correlation is −0.428. We take the weaker correlation as the more honest number. “Exposure to clinical labels” is a three-level summary of training regimes that differ in which archive was used, how it was filtered, and whether labels entered as fitting targets or only as thresholds, and twenty-two tools span far more of that heterogeneity than five. A perfectly monotone gradient over twenty-two tools would be more surprising than a partial one, and the five-predictor monotonicity should be read as a property of a small and deliberately spread sample rather than as the stronger result.

*The headroom confound does not survive the larger panel* (Fig 5b). Across five predictors, change and global AUROC were perfectly rank-correlated and the two explanations could not be separated. Across these twenty-two the rank correlation between global AUROC and change is −0.256 (p = 0.25), and −0.341 (p = 0.25) among the thirteen high-confidence representatives. Whatever orders these changes, it is not overall accuracy.

*The gradient holds, and strengthens once headroom is removed.* Over the thirteen high-confidence representatives the raw correlation between exposure and change is −0.538 (exact permutation p = 0.060); across all twenty-two it is −0.428 (p = 0.049). Against the degradation curve fitted on this same intersection, the correlation between exposure and the residual is −0.821 (p = 0.0016) and −0.658 (p = 0.0014) respectively. We note that the raw correlation on the pre-registered primary set sits at the conventional threshold rather than clearly below it, and we do not overstate it; the headroom-corrected correlation and the gene-level contrast below are the firmer evidence.

*The same hypothesis, tested on genes rather than predictors.* The quantity that matters is not which group predicts better, the supervised group does, by 0.0259 AUROC even within genes, but how much each group *loses* when gene identity is removed. Contrasting the never-trained and supervised groups globally and again inside genes gives a difference that can be resampled over genes rather than over predictors (Table 3): The contrast is drawn in Fig 5c.

**Table 3.**
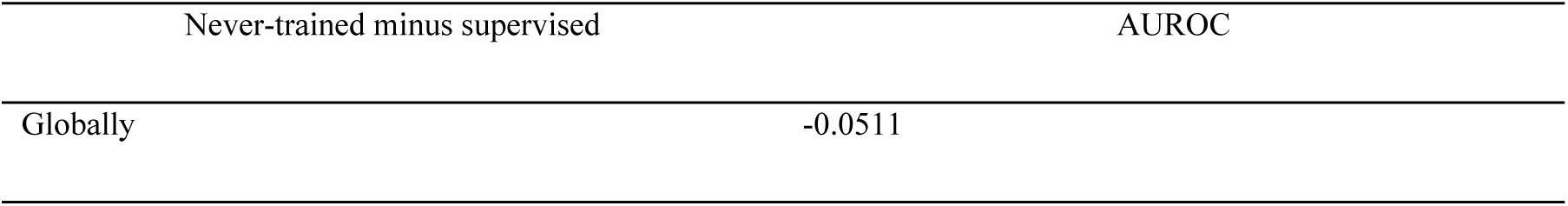
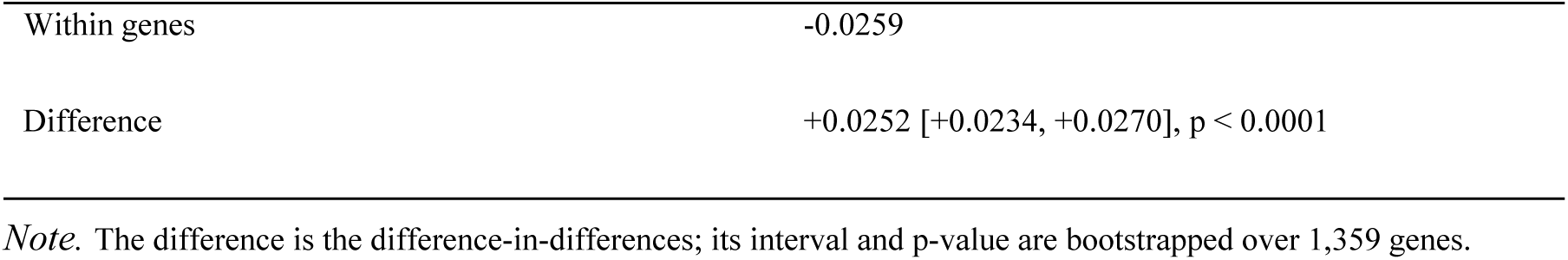
Never-trained minus supervised AUROC, globally and within genes.

Predictors never exposed to clinical labels sit 0.0511 AUROC behind the supervised ones on a conventional benchmark. Inside genes they are 0.0259 behind. Roughly half the supervised advantage is gene-level information rather than variant-level discrimination, bootstrapped over 1,359 genes with an interval well clear of zero. This is the strongest form of the claim in this study, and the only one whose sample size is genes rather than tools.

Three predictors gain under gene control, and all three are level 0: MisFit_S (+0.0502), MPC (+0.0550) and popEVE (+0.0080). MisFit_S estimates selection coefficients from allele counts in 236,017 individuals [24], MPC measures regional missense constraint from population sequencing, and popEVE combines an unsupervised evolutionary model with population data. None uses clinical labels, and all three are the only models in the panel whose training signal is population genetics or evolution rather than annotation.

One apparent inconsistency deserves stating rather than leaving to be found. ESM-2 zero-shot gains +0.0167 under gene control on the mitochondrial subset, while ESM1b in this panel loses 0.0275, two protein language models with opposite signs. They are different models scored on very different variant sets (133 evaluable genes against 10,458), and dbNSFP’s ESM1b column is a released per-substitution score rather than the masked-marginal quantity we compute. We do not treat the two as comparable, and neither is used as evidence for the mechanism on its own.

We state the limits. The exposure categories are our assignment rather than a randomised treatment, and the predictors differ in age, training data and architecture as well as in exposure. What we can say is that the ordering is not produced by differences in overall accuracy, because we constructed that alternative explicitly, and that the gene-level contrast is large, tightly bounded, and measured on identical data.

The mechanism is in any case not required for the central claim. Whatever produces the reversal, the reversal itself is a measured property of two independently curated benchmarks and stands on its own.

The mechanism makes a further directional prediction we can test: a predictor that never saw any clinical label should gain the most under gene control. ESM-2 zero-shot is that case. On the subset where ESM-2 scores exist (6,839 variants, 890 genes, 133 evaluable), the prediction holds (S4 Table). ESM-2 gains +0.0167 under gene control, VARITY_R_LOO +0.0043, AlphaMissense −0.0005 and gMVP −0.0130.

ESM-2 gains most, exceeding the larger supervised gain in 91.5% of 2,000 gene-level bootstrap resamples. The test is small: 133 evaluable genes against 1,860 in the main analysis, because ESM-2 was scored only on the mitochondrial and control sets. It supports the direction of the mechanism but is far too small to reproduce the reversal, which does not appear at this sample size.

*The reversal replicates on an independently curated benchmark.* Everything above rests on a variant table we assembled ourselves, so a reviewer is entitled to ask whether the reversal is a property of variant effect prediction or of our curation. We repeated the analysis on the ProteinGym [25] clinical substitutions set: 62,727 variants across 2,525 proteins, assembled by a different group through a different pipeline with its own pathogenic and benign annotation. Proteins were matched to UniProt by exact sequence identity rather than through an ID-mapping service, giving 2,188 of 2,525 proteins with 100% of wild-type residues validated against the canonical sequence; 49,260 variants in 2,046 proteins carry scores from all three predictors, of which 929 proteins are evaluable (Fig 4d; Table 4).

**Table 4.**
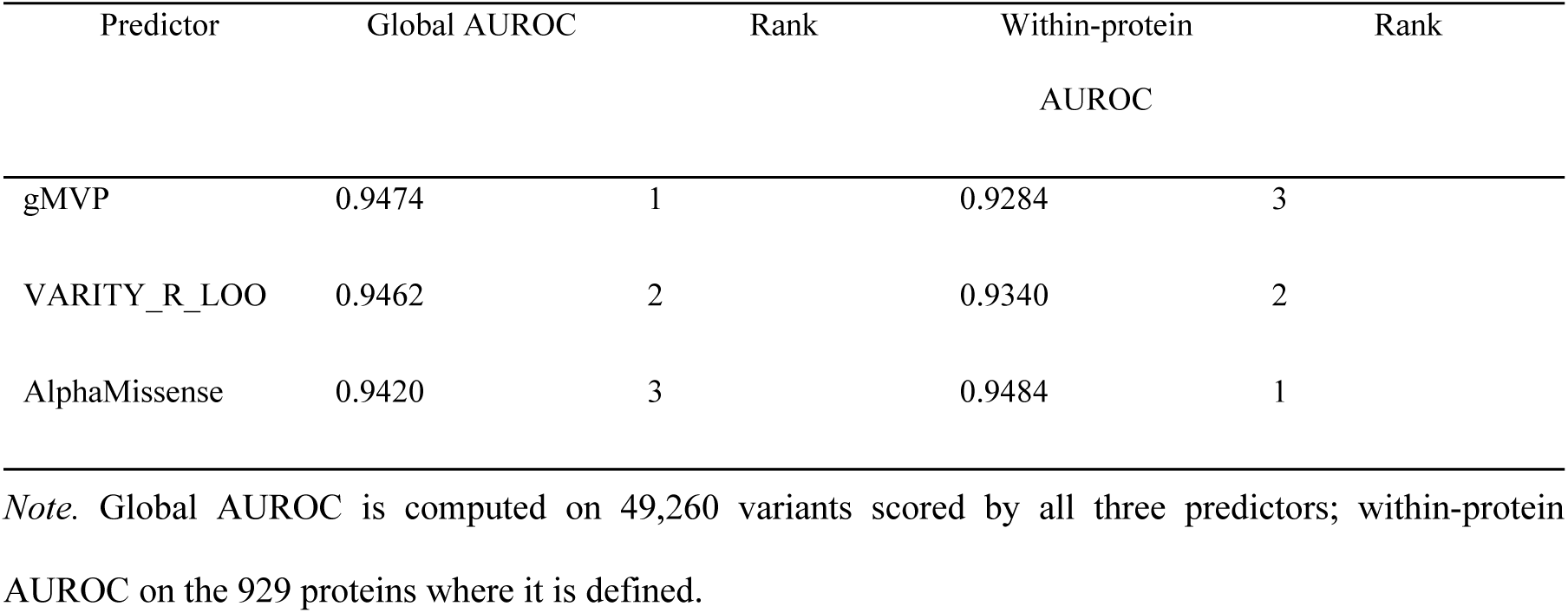
Independent replication on the ProteinGym clinical substitutions benchmark.

The ordering inverts completely, and here all three pairwise comparisons reverse sign rather than two of three: VARITY - AlphaMissense goes from +0.0042 to −0.0144 (p < 0.0001), AlphaMissense - gMVP from −0.0055 to +0.0200 (p < 0.0001), and VARITY - gMVP from −0.0012 to +0.0056 (p = 0.041). AlphaMissense again moves from last to first; gMVP moves from first to last.

Two independently assembled benchmarks, with different curators, different inclusion criteria and largely different variants, therefore produce the same inversion. The effect is a property of how ClinVar-derived benchmarks are structured, not of our particular table.

*Consequence.* The confound is therefore not a uniform inflation that cancels in comparisons. It changes the ordering. A benchmark reporting global AUROC on ClinVar can rank one predictor above another when the reverse holds for the question a clinician actually asks.

### 2.5 Experiment arbitrates between the two rankings

Both rankings are computed from the same ClinVar labels, so nothing internal to ClinVar can establish which one measures the more useful quantity. A reviewer is entitled to ask why a metric of our construction should be preferred to the conventional one, particularly since the newest predictor wins under ours.

Deep mutational scanning [26] answers this from outside. Such assays are collected in public repositories [27], are admissible as functional evidence under the ACMG/AMP framework [28], and are an established external benchmark for variant effect predictors [29]. Such an assay measures every substitution in a single protein, so agreement with one is structurally a within-protein comparison: no between-gene label structure exists to exploit and the gene-identity null is not even definable. If gene structure inflates global ClinVar AUROC, then the within-gene ClinVar ranking, not the global one, should predict which predictor agrees with experiment. That is falsifiable, and the result could have gone against us.

We took every human assay in ProteinGym, 96 of 217, and kept those whose target sequence matched a UniProt canonical sequence exactly, giving 55 assays across 48 proteins. Predictor scores were extracted per protein from the full releases rather than from our ClinVar-restricted tables, because a ClinVar-restricted join would recover only the clinically ascertained minority of each assay. Within each assay, all three predictors are correlated against experimental fitness on the substitutions all three score, so no correlation is computed on a subset the others do not share; 47 assays have at least 100 such substitutions (median 1,603, 40% of the assay). Agreement with experiment is shown in Table 5.

**Table 5.** Agreement with experimental deep mutational scanning across 47 human ProteinGym assays, against the two ClinVar rankings.

| Predictor | Mean Spearman vs<br>experiment | Rank | Global ClinVar rank | Within-gene<br>ClinVar rank |
| --- | --- | --- | --- | --- |
| AlphaMissense | 0.4170 | 1 | 3 | 1 |
| VARITY_R_LOO | 0.4132 | 2 | 2 | 2 |
| gMVP | 0.3906 | 3 | 1 | 3 |

The experimental ordering matches the within-gene ClinVar ordering exactly and inverts the global one. Because there are only three predictors, the rank correlation is ±1 by construction and carries little information; the paired tests across assays are the evidence (Fig 4e; Table 6):

**Table 6.**
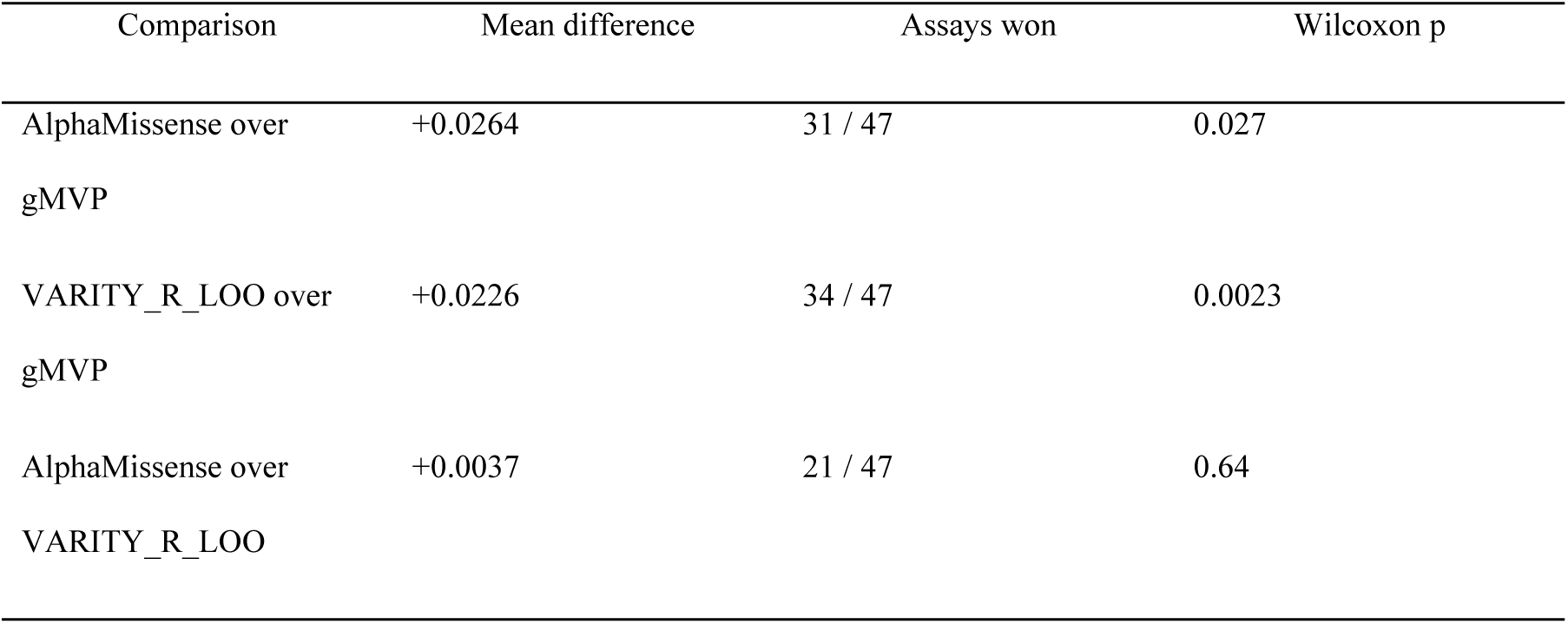
Paired comparisons of agreement with experiment across the 47 assays.

| Comparison | Mean difference | Assays won | Wilcoxon p |
| --- | --- | --- | --- |
| AlphaMissense over gMVP | +0.0264 | 31 / 47 | 0.027 |
| VARITY_R_LOO over gMVP | +0.0226 | 34 / 47 | 0.0023 |
| AlphaMissense over VARITY_R_LOO | +0.0037 | 21 / 47 | 0.64 |

Two of the three pairwise comparisons are resolved by experiment, and both favour the ordering that gene-controlled evaluation gives over the one conventional benchmarking gives. Both involve gMVP, the predictor that global ClinVar AUROC ranks first and within-gene AUROC ranks last: on experimental data it is last, significantly so against both competitors. The third comparison, AlphaMissense against VARITY, is the one within-gene evaluation resolves most weakly (a 57% win rate over genes), and experiment does not resolve it at all (p = 0.64). The two analyses agree about what is and is not decidable.

We are careful about what this does and does not show. Agreement with a functional assay is not the same quantity as clinical pathogenicity, assays differ in what they measure, and 47 assays over three predictors is not a large basis. This arbitrates between two rankings of the same three tools; it does not establish within-gene AUROC as a measure of clinical utility. But the conventional ranking is not merely unhelpful here, it is inverted relative to experiment, and that is difficult to reconcile with the view that global ClinVar AUROC is measuring variant-level discrimination.

### 2.6 The effect is not an artefact of sparsely annotated genes

The immediate objection is that genes with one or two ClinVar entries are label-pure trivially, so the null’s power might reflect sparse annotation rather than a property of the benchmark. We tested this by restricting to progressively better-sampled genes (Fig 2b; S5 Table).

The fraction of label-pure genes falls almost tenfold across this range while the null’s AUROC does not move, staying between 0.920 and 0.924. The margin available to variant-level prediction does not widen as annotation improves; it narrows, from 0.035 to 0.021. On the 649 most thoroughly characterised genes in ClinVar, comprising 90,407 variants, AlphaMissense exceeds the gene-identity null by approximately two AUROC points.

### 2.7 The pattern holds in every subcellular compartment

We partitioned the dataset by UniProt subcellular localisation and repeated the comparison within each compartment (Fig 2c; S6 Table).

The null lies between 0.835 and 0.935 in every compartment, and in none does variant-level prediction exceed it by more than 0.109. Absolute predictor performance varies across compartments by 0.028 AUROC, but so does the null, and the two are positively associated (Pearson r = 0.55 across ten compartments; given n = 10 we treat this as suggestive rather than established).

This is the result with the most direct methodological consequence. A study that benchmarks predictors within a chosen compartment and compares against the genome-wide average will observe a difference, and that difference is not attributable to the biology of the compartment without first accounting for how much of it the null explains.

### 2.8 Worked example: testing a claim of specialisation

The three recommendations above are best shown on a case where the answer was not known in advance, and we use our own earlier work. Mitochondrial disease genes have been proposed as needing specialised variant interpretation on the strength of benchmark differences from the genome-wide average. Reconstructed on MitoCarta 3.0 [30] membership alone, with a control set matched gene-by-gene on variants per gene and pathogenic fraction, that case largely dissolves: neither null model shows any mitochondrial-versus-control difference, so where a predictor deficit appears it is genuine but small (1.3 to 3.0 AUROC points, and significant for gMVP in both analysis sets, AlphaMissense only at the >=2-star floor); mitochondria-specific biophysical features contribute nothing measurable in ablation; and a specialised classifier rebuilt on corrected data reaches 0.911 against AlphaMissense’s 0.940 on the same variants. On the three mitochondrial deep mutational scanning assays in ProteinGym, all predictors retain substantial agreement with experimental fitness (Spearman 0.50 to 0.65), confirming that the signal they carry is not an artefact of ClinVar’s gene structure.

The full analysis, including the compartment comparison, the feature ablation and the assay-level correlations, is in S2 Text (S7 Table, S8 Table, S9 Table, S10 Table).

### 2.9 Calibration and protein-level uncertainty

AUROC measures ranking, not reliability. A predictor can rank variants well while its scores mean nothing as probabilities, and clinical use depends on the latter. We therefore measured calibration for every predictor, with temperature scaling fitted on training clusters only, and split-conformal prediction sets at a 90% target coverage (Table 7, Fig 6; Methods, “Calibration and uncertainty”).

**Fig 6.**
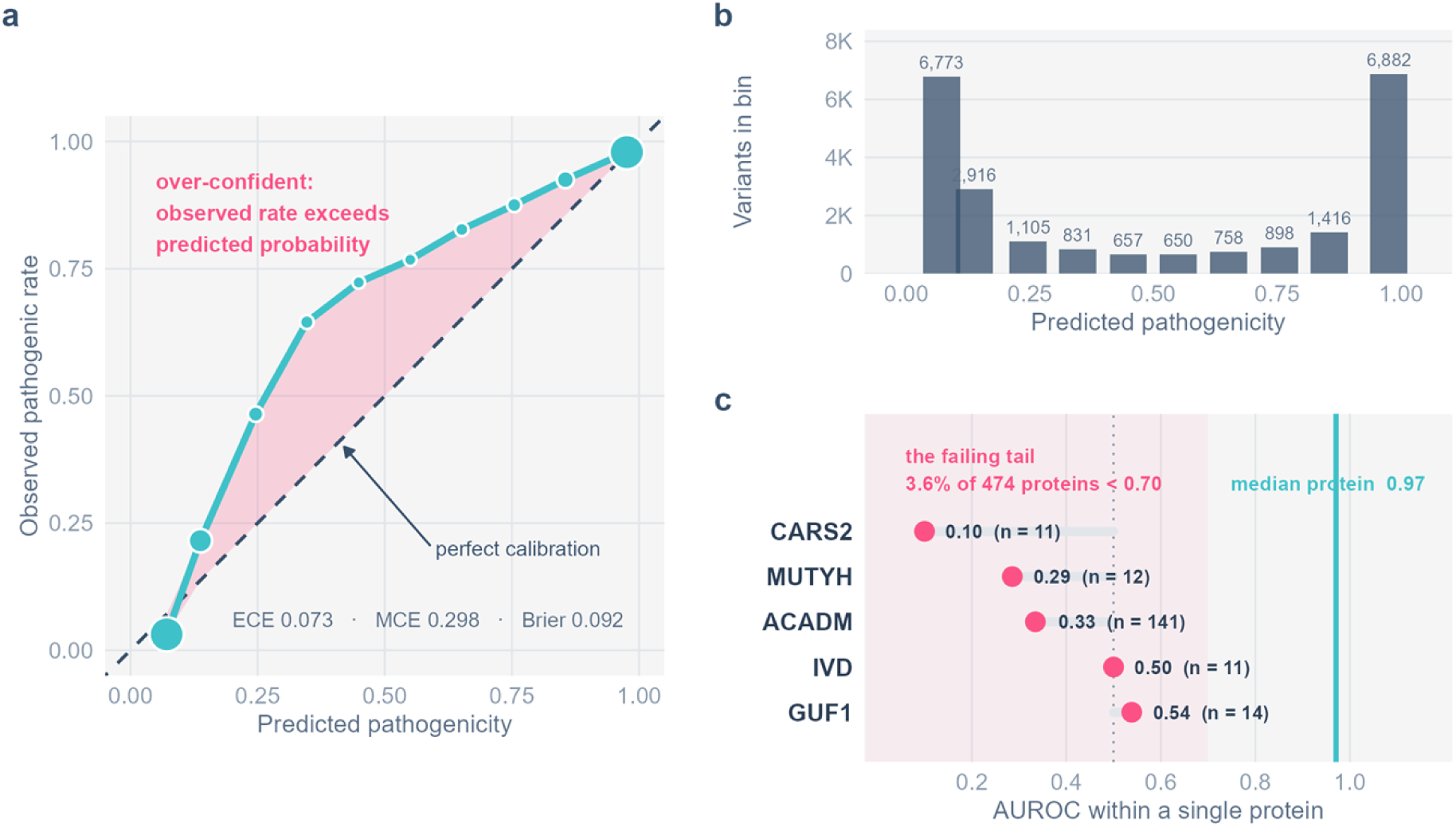
Calibration. **a**, Reliability curve with miscalibration drawn as a filled ribbon against the diagonal. **b**, Bin occupancy, showing which parts of the curve carry weight. **c**, The per-protein failing tail, named.

**Table 7.** Calibration and split-conformal prediction at 90% target coverage.

| Predictor | ECE | ECE after<br>scaling | Brier | Conformal<br>coverage | Variants<br>returned as<br>uncallable |
| --- | --- | --- | --- | --- | --- |
| VARITY_R_LOO | 0.027 | 0.009 | 0.076 | 0.900 | 0.2% |
| gMVP | 0.050 | 0.037 | 0.090 | 0.900 | 4.4% |
| AlphaMissense | 0.073 | 0.070 | 0.092 | 0.900 | 6.5% |
| ESM-2 zero-shot | 0.267 | 0.186 | 0.239 | 0.900 | 37.4% |

Calibration separates the predictors in a way AUROC does not. AlphaMissense and VARITY differ by 0.007 AUROC on the common intersection but by a factor of approaching three in expected calibration error, and AlphaMissense barely improves under temperature scaling while VARITY’s error falls to 0.009. ESM-2’s raw output is a log-likelihood ratio rather than a probability and is correspondingly uncalibrated; the conformal analysis makes the consequence concrete, returning both labels, an explicit refusal to call, for 37% of variants against 0.2% for VARITY.

Conformal coverage lands on the 0.900 target for every predictor, which is the guarantee the method provides and a form of uncertainty that does not depend on the scores being calibrated in the first place.

At the protein level, restricting to proteins with at least ten annotated variants and both classes present, median per-protein AUROC is 0.971 for AlphaMissense and VARITY, 0.944 for gMVP and 0.973 for ESM-2. Between 3.6% and 6.4% of proteins fall below 0.70 for a given predictor. These are the proteins on which computational evidence should carry reduced weight, and they are listed per predictor in the deposited result tables.

## 3. Discussion

The central finding is easily stated. When a missense pathogenicity predictor is benchmarked on ClinVar with a random variant-level split, roughly 0.92 of the 0.96 AUROC it reports is achievable without looking at the variant at all. This does not mean predictors are uninformative; it means the quantity usually reported is not a clean measure of variant-level discrimination, and that differences of a few AUROC points between tools, or between gene sets, sit inside the range that gene-level label structure can produce on its own.

*Implications for evaluation.* We suggest three changes. First, a gene-identity null should be reported alongside any ClinVar benchmark; it costs nothing to compute and calibrates the reader’s expectation. Second, leave-gene-out or homology-controlled splitting should be the default. Under leave-gene-out the null is exactly 0.500 by construction, so any performance above chance is genuinely variant-level. Third, claims that a gene set requires specialised modelling should be supported by a comparison against a matched control set, not against the genome-wide average.

*Relation to known circularity and to gene-level heterogeneity.* Type 1 and type 2 circularity concern shared variants and shared proteins between training and test data [12]. The confound described here survives the elimination of both: no variant and no protein need be shared for gene-level label structure to inflate a score, because the structure is a property of how the archive was assembled rather than of any particular split.

It is also distinct from the gene-level calibration problem documented recently [13, 14]. Tejura and colleagues show that score intervals calibrated genome-wide do not hold within individual genes, so a variant can receive materially wrong evidence strength depending on which gene it sits in. We do not dispute or duplicate that result; our compartment analysis is consistent with it, and it is the stronger finding for immediate clinical practice.

The distinction is between calibration and discrimination. Their work asks whether a score means the same thing everywhere. Ours asks how much of the reported ranking performance survives removing the variant from consideration. A predictor could be perfectly calibrated in every gene and still owe most of its AUROC to gene identity. The remedies diverge accordingly: gene-level miscalibration is addressed by calibrating per gene, whereas near-sufficiency of gene identity is addressed only by changing the split or by reporting the null alongside the score.

Our leave-one-out result shows the distinction is not academic. The field’s existing correction for training-set contamination, applied exactly as its authors intended, removes 0.002 of an effect worth 0.036 to 0.044, because it operates on variants when the structure it needs to remove operates on genes.

*What the two strongest tests add.* Both rankings above derive from the same ClinVar labels, so ClinVar cannot adjudicate between them. Two lines of evidence external to that circle do. Agreement with 47 deep mutational scanning assays, data with no between-gene label structure to exploit, and therefore no gene prior to inflate anything, reproduces the within-gene ordering and inverts the conventional one, with both resolvable comparisons favouring gene control. And across twenty-two predictors scored on identical variants, the never-trained group closes half of its global deficit once gene identity is removed (+0.0252 [+0.0234, +0.0270] over 1,359 genes). The first says the gene-controlled ranking is the one that predicts experiment; the second says the advantage conventional benchmarks award to clinically supervised models is about half gene-level bookkeeping.

Neither result depends on our null model being a sensible predictor, and neither requires the exposure classification to be correct, the difference-in-differences would survive any relabelling that keeps the never-trained and supervised groups distinct.

*On specialisation.* We came to this question from the opposite direction, having attempted to build a mitochondria-specific predictor. That work failed for instructive reasons: the gene set was contaminated by a phenotype-keyword filter, the mitochondria-specific features contributed nothing measurable in ablation, and a general-purpose predictor outperformed the specialised one. Reconstructed properly, the case for mitochondrial specialisation largely dissolves, though not entirely, since a small real deficit remains at high annotation confidence, and we report it as such.

*“But the null is not a usable predictor.”* This is the strongest objection to our framing and we want to meet it directly rather than list it as a limitation. It is entirely correct: the gene-identity null collapses to exactly 0.500 under leave-gene-out splitting, cannot score a variant in a gene it has not seen, and would be useless in a clinic. AlphaMissense generalises to genes with no ClinVar entries at all; the null cannot.

That is precisely the point. We do not propose the null as a predictor, and no part of our argument requires it to be one. We propose it as a *measuring instrument* for benchmarks. Its uselessness in the clinic is what makes it diagnostic: if a quantity that cannot generalise to a new gene reproduces 0.921 of a benchmark score, then that benchmark is substantially measuring something which does not generalise to a new gene either. The comparison is not “use the null instead,” it is “know how much of your reported number the null already accounts for.”

The within-gene analysis makes this concrete. There the null is not merely useless but undefined, the gene prior is constant within any single gene, and the predictor ordering changes. That is not a statement about the null’s clinical value. It is a statement about what global ClinVar AUROC ranks.

*Other limitations.* Within-gene evaluation requires genes carrying at least three variants of each class, which are the better-characterised genes and are not representative of ClinVar as a whole. The three-predictor reversal is established on 1,860 such genes and the dbNSFP contrast on 10,458, but neither is demonstrated for genes with sparse annotation. Our compartment analysis assigns each protein a single primary localisation, which is a simplification. The correlation between predictor and null performance across compartments rests on ten points. Experimental validation rests on 47 deep mutational scanning assays for the arbitration and three for the mitochondrial comparison, all of which are necessarily restricted to proteins that have been assayed at scale and are not a random sample of the proteome. And AlphaMissense is calibrated on ClinVar, so its performance here is not independent of the archive whose structure we are characterising, a point that applies to any supervised predictor evaluated this way.

### 3.1 Conclusions

When a missense pathogenicity predictor is benchmarked on ClinVar with a random variant-level split, a model that knows only which gene each variant sits in reaches AUROC 0.921. Four current predictors exceed this by 0.036 to 0.044. The margin does not widen as gene annotation improves, it narrows, to 0.021 on the best-characterised genes, and it holds in every subcellular compartment we examined. Leave-one-out correction, the standard remedy for training-set contamination, addresses two thousandths of it. Across twenty-two predictors on identical variants, roughly half the advantage that conventional benchmarks give clinically supervised models disappears once gene identity is removed.

None of this means predictors are uninformative. On deep mutational scanning assays, where no gene-level structure exists to exploit, predictors retain substantial agreement with experimental fitness, and across 47 such assays that agreement follows the gene-controlled ranking rather than the conventional one. What it means is that the quantity conventionally reported is not a clean measure of variant-level discrimination, and that differences of a few AUROC points, between tools, or between gene sets, fall inside the range gene-level label structure produces on its own.

Three changes follow. Report a gene-level null alongside any ClinVar benchmark; it costs one function call and we release the implementation. Make leave-gene-out or homology-controlled splitting the default, under which the null is exactly 0.5 and everything above chance is genuinely variant-level. And test claims that a gene set needs specialised modelling against a matched control set rather than the genome-wide average. Our own mitochondrial case study is the cautionary example: the apparent case for specialisation dissolved under each of these controls in turn.

## 4. Materials and methods

### 4.1 Variant dataset

ClinVar variant_summary.txt.gz (release recorded in data/rebuild/build_metadata.json with MD5) was filtered to GRCh38 assembly rows with a gene symbol and an unambiguous classification: pathogenic or likely pathogenic without any benign or conflicting term, or benign or likely benign without any pathogenic or conflicting term. Protein consequences were parsed from the HGVS Name field. Synonymous substitutions were excluded.

Each variant was mapped to a UniProt reviewed human accession by gene symbol and retained only if the wild-type residue matched the canonical sequence at the stated position (93.1% of gene-mapped variants). Duplicate protein-level substitutions were collapsed, retaining the highest review-status rating. ClinVar review status was converted to a star rating; analyses are reported at both the full set and a ≥2-star floor, declared in advance as the primary analysis set.

### 4.2 Mitochondrial and control sets

Mitochondrial membership was defined solely by presence in MitoCarta 3.0, matching on primary symbols and listed synonyms. No phenotype or disease keywords were used at any stage. The control set was drawn from non-MitoCarta genes, binned by variants per gene and pathogenic fraction, with one control gene sampled per mitochondrial gene from the matching bin (seed 42).

### 4.3 Homology control and splitting

Proteins carrying a validated variant were clustered with MMseqs2 [31] easy-cluster at -- min-seq-id 0.3 -c 0.8 --cov-mode 0. Cluster assignment is the grouping variable for leave-cluster-out cross-validation, so no two proteins above the identity threshold are split across a train/test boundary. Version hash and full command are recorded in data/rebuild/cluster_metadata.json.

### 4.4 Null models

The in-sample gene prior scores each variant by the pathogenic fraction of its own gene computed over the whole dataset. The random-split variant recomputes this on each training fold of a stratified 10-fold split. The leave-gene-out and leave-cluster-out variants recompute it on training folds from which the test gene or cluster is absent; unseen genes receive a single global constant of 0.5. Using a per-fold training base rate instead induces a spurious ranking across held-out folds and must be avoided.

### 4.5 Comparator predictors

AlphaMissense scores come from the bulk amino-acid substitution release, joined on UniProt accession and protein-level substitution so that no genomic liftover is required (98.7% coverage). VARITY scores come from the precomputed prediction archive, which is also keyed by UniProt accession and so joins directly. gMVP scores come from the precomputed release for all canonical-transcript missense variants, which is keyed by gene symbol and Ensembl transcript position.

The gMVP join therefore carries a risk worth stating: Ensembl and UniProt canonical transcripts do not always agree, and where they differ the same position number denotes a different residue. Every join here requires the source’s reference residue to equal our wild-type residue, which makes it self-validating, a transcript mismatch almost always produces a reference-residue mismatch, and the variant is dropped rather than silently mis-scored. For gMVP this yielded 179,238 of 197,904 variants (90.6%), with 4,256 dropped on reference-residue mismatch; that count is reported separately rather than folded into the coverage figure.

*Circularity in the supervised comparators.* VARITY is trained on ClinVar, so scoring our ClinVar variants with the standard VARITY_R output would report its performance on its own training data. The release ships VARITY_R_LOO, a leave-one-out score in which each variant is predicted by a model that did not see it; that is the column we use. Both are retained, and the gap between them is itself a direct measurement of the circularity that motivates this study.

ESM-2 zero-shot scores use masked-marginal scoring [32] with esm2_t33_650M_UR50D [22]: the variant position is masked, and the score is log p(mut) - log p(wt) at that position. Proteins longer than the model’s 1,022-residue limit are scored in a window centred on the variant, never by N-terminal truncation.

### 4.6 Within-gene evaluation and its two intersections

Within-gene AUROC is computed independently in each gene carrying at least three variants of each label class, then summarised across genes. Genes with only one label class are excluded because the quantity is undefined there.

Two intersections are reported and they are not interchangeable. The three modern predictors (AlphaMissense, VARITY, gMVP) each cover most of ClinVar, giving 169,989 variants and 1,860 evaluable genes; this is where the ranking reversal is established. PolyPhen-2 and SIFT are supplied through the EBI Proteins API for annotated natural variants only, so requiring all five predictors reduces the intersection to 59,045 variants and 1,055 evaluable genes. The five-predictor set is used only for the ClinVar-exposure comparison, where spanning two decades of methodology matters more than sample size.

The two intersections are computed after the predictor set is fixed, not before, otherwise restricting to a subset would silently inherit the intersection of all five. Results are written to separate files (within_gene_ranking_modern.* and within_gene_ranking_all.*) so that neither analysis overwrites the other.

Because roughly a third of evaluable genes are perfectly separated by at least one predictor, the mean is reported alongside the median, a variant-weighted mean, and paired Wilcoxon signed-rank tests [33], and the ordering is re-checked with ceiling genes excluded. Per-gene “win” counts exclude genes where two or more predictors tie, since assigning ties by column order produces an ordering that depends on the order chosen.

### 4.7 Headroom control

The change from global to within-gene AUROC is confounded with global AUROC itself: a weaker predictor has more room to move. To construct the null in which that is the only mechanism, each predictor’s score was converted to within-column percentile ranks and mixed with uniform noise, s’ = (1 - a)·rank(s) + a·u, over twenty levels of a from 0 to 0.95, with five independent noise draws averaged at each level. Mixing is applied to the whole score column at once, so between-gene and within-gene discrimination degrade together, as they would for a genuinely weaker predictor; and it is monotone in the original ordering at a = 0.

Global and within-gene AUROC were recomputed at every level, giving 95 degraded points across the five predictors. The same construction is applied independently to the dbNSFP panel on its own intersection, with nineteen levels and three noise draws, giving 418 degraded points across twenty-two predictors; the curve used for those residuals is fitted to that sweep, not to the five-predictor one. A quadratic in global AUROC was fitted to those points, excluding the undegraded ones, so that the observations under test do not help define the curve they are tested against, and each predictor’s residual is its observed change minus the change that curve predicts at its own global AUROC. Residual confidence intervals come from a 2,000-replicate bootstrap resampling whole genes, with the fitted curve held fixed.

Within-gene AUROC in this analysis is computed through the Mann-Whitney identity rather than by looping a general AUROC routine over genes, which the sweep would otherwise make prohibitively slow; the two agree to within 10⁻⁹ and the check is asserted at runtime.

### 4.8 Rebuilding the specialised classifier

The mitochondria-specific classifier from our earlier work reported AUROC 0.890, computed on the contaminated variant set. That number is not recomputable here and is not comparable with values measured on the corrected data, so it is not used. The pipeline was instead rebuilt: the interpretable feature groups above, engineered from the UniProt canonical sequence, together with the ESM-2 zero-shot masked-marginal score, fitted with L2-regularised logistic regression under leave-cluster-out cross-validation on the ≥2-star mitochondrial set. The original used 128 principal components of ESM-2 embeddings rather than the zero-shot score, and split randomly rather than by homology cluster; both differences make the rebuilt figure a reconstruction of the design rather than a reproduction of the original result, and both are stated wherever the number appears.

### 4.9 The dbNSFP panel

All 37 per-predictor score columns were extracted from dbNSFP 5.3.1a [23] and joined to our variants on UniProt accession, protein position and both residues. dbNSFP stores accession and position as parallel semicolon-separated lists across alternative transcripts; a row matches if any pair matches ours and dbNSFP’s own reference residue equals our wild-type residue, which makes the join self-validating against transcript mismatch in the same way as the other comparators. 195,164 of 197,904 variants (98.6%) carry at least one score.

Seven columns encode damage as a low value (SIFT, SIFT4G, PROVEAN, FATHMM, ESM1b, popEVE, LRT) and are negated before use. Polarity is taken from dbNSFP’s documented convention rather than from whichever direction scores above 0.5, which would fit the encoding to the labels; the analysis stops if any retained predictor remains inverted afterwards.

Exposure to clinical labels was classified from each tool’s published training description and fixed before any score was extracted (S1 Text). Two exclusions follow from the panel’s structure rather than from any result. Ensembles that take other listed predictors as input features are not statistically independent of them and are excluded: MetaSVM, MetaLR, MetaRNN, M-CAP, REVEL, BayesDel (both variants) and ClinPred. Where one model contributes several columns, one pre-specified representative is used, so that VARITY, which ships as four columns, is not counted four times.

Predictors covering fewer than half our variants are dropped, a threshold set in advance. Every predictor in the release clears it once per-transcript values are parsed correctly: dbNSFP stores one entry per transcript and the first is often ‘.’, so reading element zero silently discards a real score and had put coverage near 50% across the whole panel. Twenty-two representatives remain, and all reported values are computed on their common intersection of 112,248 variants in 10,458 genes. Scoring each predictor on its own coverage subset instead leaves every sign unchanged and shifts the changes by at most 0.0105 (mean 0.0038), so the common intersection is used on principle rather than because it rescues the result. On the earlier extract, where a per-transcript parsing error had suppressed coverage to roughly 50%, the two did disagree in sign for several predictors, a reminder that comparisons across differing subsets are unsafe even when they happen to agree.

Correlations between exposure and change use exact permutation p-values over all orderings, because thirteen predictors is too few for the asymptotic approximation to be trustworthy. The difference-in-differences contrasts the never-trained and supervised groups globally and again within genes, and is resampled over the 1,359 genes evaluable for every predictor in both groups, with 500 replicates.

### 4.10 Statistics

Discrimination is reported as the area under the receiver operating characteristic curve [34]. Confidence intervals come from a bootstrap [35] resampling whole clusters (or whole genes, for the all-ClinVar analyses) with replacement, 2,000 replicates for cluster-level and 500 for gene-level analyses. Comparisons between independent sets resample each side separately and report the distribution of the difference, with a two-sided bootstrap p-value against a null of zero difference.

### 4.11 Calibration and uncertainty

Expected calibration error [36] uses ten equal-width bins, alongside the Brier score [37]. Temperature scaling [38] fits a single temperature on the logit scale using training clusters only and is applied to held-out clusters; being monotone it cannot change AUROC, only reliability, which is the point, ranking and calibration are separate claims. Split-conformal prediction sets [39, 40] use nonconformity 1 - p(true label) with the calibration quantile taken on training clusters at a 10% target error rate. A set containing both labels is an explicit refusal to call the variant rather than a forced binary decision. ESM-2 log-ratios are mapped through a sigmoid before calibration so that the quantity is defined; this is monotone and changes no ranking.

Protein-level analysis is restricted to proteins with at least ten annotated variants and both label classes present.

### 4.12 Software

Analyses were run under Python 3.14 with numpy 2.4.3 [41], pandas 3.0.1 [42], scikit-learn 1.8.0 [43], scipy 1.17.1 [44], pyarrow 23.0.1, Biopython 1.87 [45] (pairwise alignment for ProteinGym position mapping), PyTorch 2.11.0 with CUDA 12.6 [46], and transformers 5.13.0 [47] (ESM-2 inference on an NVIDIA RTX 4060). Protein clustering used MMseqs2; the exact version hash and command line are recorded in data/rebuild/cluster_metadata.json. Figures were drawn in R [48] with ggplot2 [49] and patchwork [50]. Random seeds are fixed at 42 throughout and stated in each script.

The gene-identity null is released as genenull, a single-file module with no dependencies beyond numpy, pandas and scikit-learn. It computes the null under all three split schemes and reports a predictor’s margin over it:

python from genenull import benchmark_report report = benchmark_report(df, score=’my_predictor’, gene=’gene_symbol’)

It also reports within-gene AUROC, the gene-controlled measure under which the predictor ranking in this study inverts, and exposes the per-gene table via within_gene_auroc. A command-line entry point (python −m genenull TABLE) gives the same output for a parquet or CSV file.

It reproduces every null value in this manuscript from data/rebuild/variants_all.parquet. We release it because the barrier to reporting this baseline should be one function call.

### 4.13 Data Availability Statement

The data and source code supporting the findings of this study are openly available in the GitHub repository at https://github.com/HarriziSaad/genenull. The repository includes the genenull implementation, analysis scripts, processed datasets, and metadata required to reproduce the reported results. All external datasets used in the study are publicly available from the sources cited in the Methods.

## Acknowledgments

We thank the members of the Laboratory of Health, Environment and Biotechnology for helpful discussions and feedback on early versions of this work. We are grateful to the Centre National pour la Recherche Scientifique et Technique (CNRST) for support provided within the framework of the “PhD-Associate Scholarship - PASS” Program.

## Supporting information

*S1 Text.* The pre-registered classification of every dbNSFP predictor by its exposure to clinical labels, with the basis and reference for each, the model-family assignments, the score-polarity record, and the analysis plan. Dated amendments record every change made after the file was first written and whether it preceded any result.

*S2 Text.* The mitochondrial case study in full: predictor and null performance against a matched control set, feature ablation under leave-cluster-out splitting, the rebuilt specialised classifier, and agreement with three mitochondrial deep mutational scanning assays.

*S1 Table.* All comparators against the gene-identity null on the common intersection.

*S2 Table.* Pairwise differences between predictors, global and within-gene, with bootstrap intervals.

*S3 Table.* Per-gene win rates and Wilcoxon signed-rank tests.

*S4 Table.* ESM-2 zero-shot against the supervised predictors under gene control.

*S5 Table.* Sensitivity to the minimum number of variants per gene.

*S6 Table.* Predictor and null by subcellular compartment.

*S7 Table.* Mitochondrial minus matched control for every method and both nulls.

*S8 Table.* Margin over the gene-identity null on the mitochondrial set.

*S9 Table.* Feature ablation on the ≥2-star mitochondrial set.

*S10 Table.* Agreement with deep mutational scanning assays.

*S11 Table.* Change from global to within-gene AUROC across five predictors on the five-predictor intersection.

*S12 Table.* Observed change against the change expected from reduced global performance alone, with residuals.

## Notes

### Competing Interest Statement

The authors have declared no competing interest.

